# *Xanthomonas imtechensis* sp. nov. - a novel member of non-pathogenic *Xanthomonas* with bioprotection function from healthy rice seeds

**DOI:** 10.64898/2026.02.15.705894

**Authors:** Anushika Sharma, Prabhu B. Patil

## Abstract

Non-pathogenic *Xanthomonas* (NPX) from a diverse plant hosts are being reported on an increasing basis. There are also reports of multiple species forming communities on a single host plant, such as rice, and, given their role as core endophytes in protecting plants from pathogens, it is essential to isolate and characterization of more NPX species from diverse host plants. Using phylogenomic analysis of publicly available *Xanthomonas* genome sequences, we identified a novel clade comprising NPX strains from diverse hosts. One of the strains previously reported from our lab is from healthy rice seeds and was reported to be non-pathogenic, with bio-protection function against the bacterial leaf blight pathogen. Genomic investigation confirmed the lack of type III secretion system and its effectors, consistent with their non-pathogenic nature. These strains also harbour core and unique biosynthetic loci identified in other non-pathogenic *Xanthomonas* (NPX) strains. Further investigation using multiple genomic-based taxonomic indices indicates that these strains represent a potential new species. Hence, we propose *Xanthomonas imtechensis* sp. nov. as a new species of the genus *Xanthomonas*, with the type strain being PPL568 = MTCC 13186 = CFBP 9040 = ICMP 24395.

## Introduction

*Xanthomonas* is a well known bacterial genus associated with plants. For the last century, members of the genus have primarily been known for their pathogenic lifestyles infecting more than 400 diverse plant species (1). There are currently 39 species reported in the genus (2). However, over recent decades, growing evidence suggests existence of a world of large and diverse species with non-pathogenic lifestyles. For example, for the past 150 years, *X. oryzae* has been the only known pathogenic species in rice. However, in just 10 years, reports of four non-pathogenic *Xanthomonas* (NPX) species have emerged (3-5). It is expected that, as with pathogenic species, all plants host multiple NPX species, as in rice. Hence, it is essential to explore the diverse world of NPX species in rice and other plant hosts. This will not only help understand pathogenic lifestyles but also exploit them as microbial biocontrol agents, as the NPX species can effectively protect against pathogenic species, as reported in the case of rice (6). The advent of the genomic era and genome-based taxonomic criteria is allowing for rapid and robust identification and reporting of NPX species. Many of the novel species are closely related and exist as community-like NPX species in rice and other plants. Genome-based phylogenetic trees and also multiple genome-based taxonomic criteria like Average Nucleotide Identity (ANI), digital DNA DNA hybridization (dDDH), Alignment Fraction (AF), (7, 8) etc, are allowing identification of closely related species existing as a community and that escape the taxonomists when only one criteria like ANI is used (9). The development of tools like TYGS (Type (Strain) Genome Server) has further enabled microbiologists to identify species and even subspecies using more robust dDDH trees and cutoffs (10). Even better ANI calculations, such as orthoANI and gANI, are enabling easier identification of closely related new species (11, 12). Another advantage of genome-based studies is the added evidence for NPX status, reflected by the complete absence of the Type 3 secretion system apparatus and effectors (6, 13). At the same time, genome scans and the identification of antimicrobial loci reflect lifestyle and biocontrol function (6, 14).

While there are several reports of novel NPX species, there are also numerous non-pathogenic strains from healthy plants reported in the literature that await proper taxonomic classification or have been misclassified. This study identifies a new lineage of non-pathogenic *Xanthomonas* (NPX) strains from clade IB, originating from diverse hosts, based on a genome-based phylogenomic tree. Further genome scans confirmed the absence of T3SS, its effectors, and antimicrobial loci characteristic of NPX species and its lifestyle. Interestingly, a member of this clade from rice is reported to be non-pathogenic and to protect the host against the leaf blight pathogen. This lineage is closely related to NPX species reported in rice, such as *X. indica* and *X. sontii*. Further genome-based taxonomic investigation using multiple criteria revealed that member of the clade belongs to a novel species *Xanthomonas*.

## Material method

### Phylo–taxogenomics analysis

Representative *Xanthomonas* genomes were downloaded from the NCBI GenBank database for phylogenetic analysis and annotated using Prokka v1.14.6. (15). Roary v3.13.0 was used to generate the core gene alignment (16). Phylogenetic reconstruction was performed using the core gene alignment in RAxML v8.2.12 with the GTRGAMMA model and 1,000 bootstrap replicates (17).

Genome sequences of the study strains were retrieved from NCBI GenBank. Average Nucleotide identity (ANI) values were determined using oANI (11) with USEARCH v11.0.667 (18), and validated with ANIb (ANI BLAST) via JSpeciesWS (19). Digital DNA-DNA hybridization (dDDH) estimates were obtained with Formula 2 of the Genome-to-Genome Distance Calculator v3.0. (20). Novel strains genomes were analyzed via the Type (strain) Genome Server (TYGS), generating whole-genome phylogenies using Genome BLAST

Distance Phylogeny (GBDP) with d4 formula distances (10, 21, 22), inferred using FastME v2.1.6.1 (21).

### Pangenome Analysis

Core and accessory gene content across the novel *Xanthomonas* strain and the reference strains were determined from a Roary pangenome analysis, with gene family counts extracted for flower plot visualization using custom R scripts in RStudio (22). Functional annotation and COG classification of PPL568 were performed using eggNOG-mapper (Galaxy v2.1.8+galaxy4) (23).

### Type III Secretion System (T3SS) and Associated Effectors

T3SS gene sequences were retrieved from the genome of the Xoo strain PXO99A. Type III effector (T3E) repertoires were downloaded from the *Xanthomonas* resource database. These protein sequences were then used as queries in tBLASTn v2.12.0+ searches to identify homologs in the analysed genomes, applying thresholds of 40% amino acid identity and 60% query coverage. Presence-absence patterns of T3SS genes and T3Es were visualized as heatmaps generated in TBtools v1.108. Using MacSyFinder v2.1.3 and the TXSScan model, Secretion systems and associated factors in the analyzed strains were predicted (24).

### Genome mining and Comparative Cluster Analysis

Genome mining with antiSMASH v8.0.4 (Antibiotic and Secondary Metabolite Analysis Shell) uncovered the presence of secondary metabolite biosynthetic gene clusters in PPL568 (25). A comparative genomic analysis was performed using X-T4SS genes from *X. sontii* and *X. indica* as query sequences to identify potential homologs. The resulting genetic architectures were visualised and compared using Clinker v0.0.32 (26), which generated cluster diagrams from BLASTp alignments to assess synteny and sequence conservation.

### Biochemical profiling

The biochemical profiling of strain PPL568 was performed using the BIOLOG GEN III microplate system (Biolog Inc., USA) according to the manufacturer’s Protocol A. A 100µl aliquot of bacterial culture was dispensed into each well of the BIOLOG plate, followed by incubation at 28°C. To confirm repeatability, three independent assays were conducted on separate days. Readings were recorded at 24, 48, and 72 hours of incubation using a MicroStation 2 reader, and data were analysed with MicroLog 3/5 2.0.1 software. Only clear positive or negative (±) reactions were considered valid for analysis. The results at 48 hours were the most consistent among the recorded time points.

## Results and discussion

### Phylo-taxogenomics analysis reveals a novel lineage of NPX strains from diverse plant hosts

*Xanthomonas* species are distributed in two clades, I and II. Clade I primarily consists of NPX species, while clade II mainly consists of pathogenic species (6). Within clade I, we previously reported that sub-clade IB consists of *X. sacchari, X. sontii, X. protegens*, and *X. indica* (9). When we inspected a core-gene-based phylogenomic tree, we found a novel lineage within clade IB, consisting of the NPX strain PPL568, along with F10, LMG8989, CFBP8445, and NCPPB1131 **(Fig. 1)**. While PPL568 was earlier reported from healthy rice seeds, F10 and LMG8989 were isolated from orange, and CFBP8445 and NCPPB1131 from banana.

**Figure 1.**
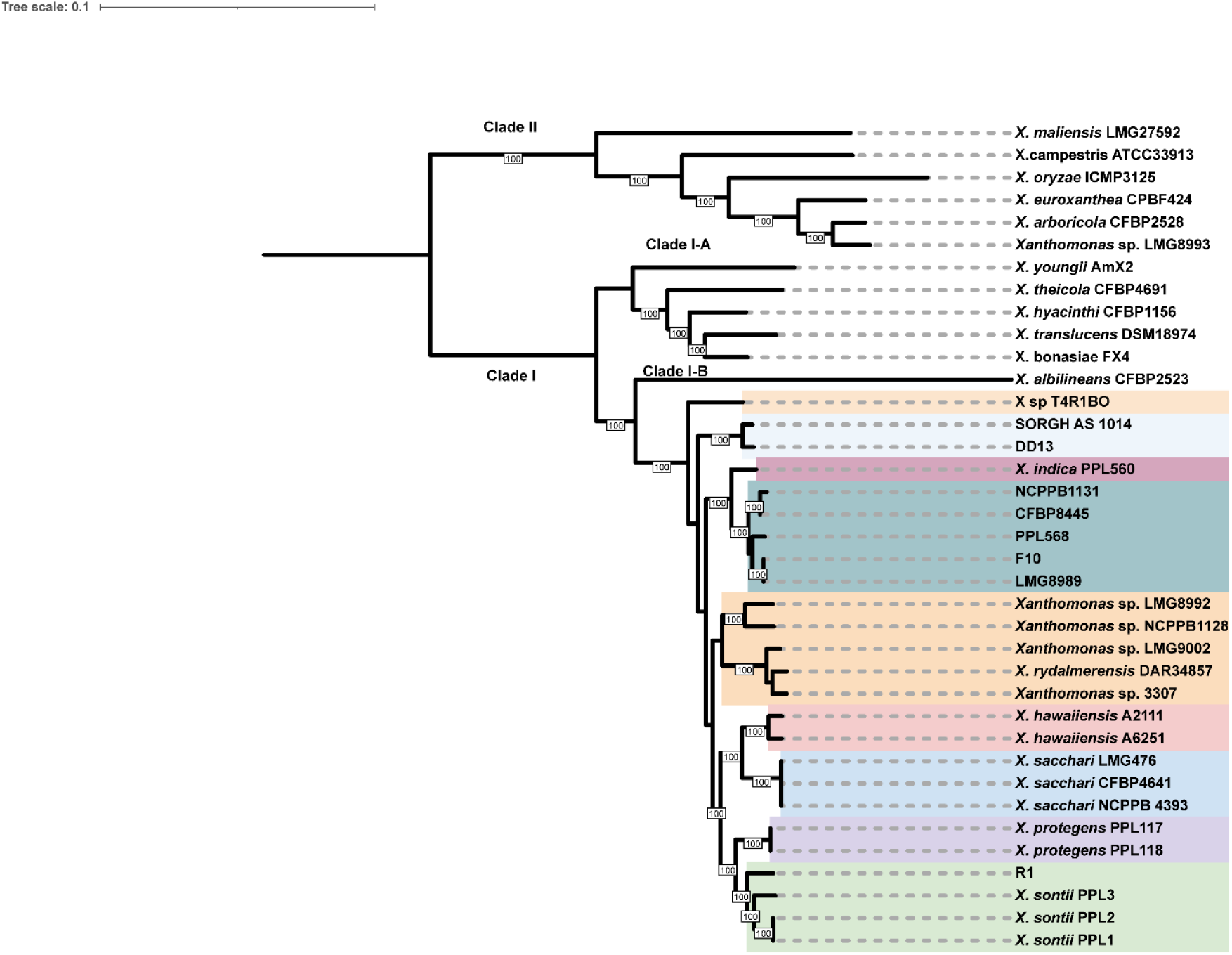
A mid-rooted maximum likelihood phylogeny based on core genes highlights PPL568 and the novel lineage in green. Clade I-B is distinguished in multiple colours, with bootstrap values >50% indicated on branches. The scale bar represents nucleotide substitutions per site.

The metadata and genomic features, along with the NCBI accession numbers of these strains, in **(Table 1)**. These bacterial strains were isolated from various hosts and regions: PPL568 (Rice, India), CFBP8445 (Banana, Samoa), F10 (Orange), LMG8989 (Orange, USA), and NCPPB1131 (Banana, Western Samoa). PPL568 exhibited a 4.75 Mb genome (69.40% GC), CFBP8445 4.87 Mb (69.30% GC), and F10 4.78 Mb (69.30% GC), all showing 100% genome completeness. LMG8989 4.47 Mb (69.30% GC), with 93% completeness and NCPPB1131 3.70 Mb (68% GC) with 64.93% completeness. Because strain NCPPB1131 has a reduced genome size and failed the CheckM test due to incompleteness, we removed it from further genome-based taxonomic analysis.

**Table 1.** Genome assembly, annotation, and metadata statistics, for the *Xanthomonas* strains analyzed in this study.

| Strain | Genome size (bps) | GC content (%) | Number of contigs | Genome coverage (x) | Genome completeness (%) | Coding sequences (CDSs) | Geographic location | Host | Accession number |
| --- | --- | --- | --- | --- | --- | --- | --- | --- | --- |
| PPL568 | 4,747,217 | 69.40 | 18 | 1,317 | 100 | 4,044 | Punjab, India | Rice | JAKLXA00000000 |
| CFBP8445 | 4,869,984 | 69.30 | 1 | 208 | 100 | 4,122 | Samoa | Banana | CP102594 |
| F10 | 4,776,116 | 69.30 | 6 | 314 | 100 | 4,043 | – | Orange | JACHNP00000000 |
| LMG8989 | 4,472,724 | 69.30 | 72 | 125 | 93 | 3,777 | USA | Orange | QUXI00000000 |
| NCPPB1131 | 3,700,000 | 68 | 3224 | 80 | 64.93 | 3224 | Western Samoa | Banana | AGHY00000000 |

Previously, PPL568 was classified as *Xanthomonas indica;* however, its dDDH value with the type strain is only 67.9% well below the 70% species cutoff, while OrthoANI at 96.3% is only borderline to the standard threshold. Strains F10, LMG8989, and CFBP8445 show around 68% dDDH and 96% OrthoANI with the *X. indica* type strain, PPL560^T^. Strains F10, LMG8989, and CFBP8445 show dDDH values of around 84% and OrthoANI values of 98%, with PPL568 exceeding intraspecies thresholds, forming a cohesive, novel lineage **(Table 2)**. And PPL568 shares dDDH values of approximately 50-51% and ANI values of 92-93% with representative clade IB strains from other *Xanthomonas* species—both below species delineation cutoffs **(Supplementary Table 1)**. This genomic coherence across diverse hosts supports the proposal of a new *Xanthomonas* species distinct from *X. indica*, as justified by genome-based tree and taxonomic indices.

**Table 2.**
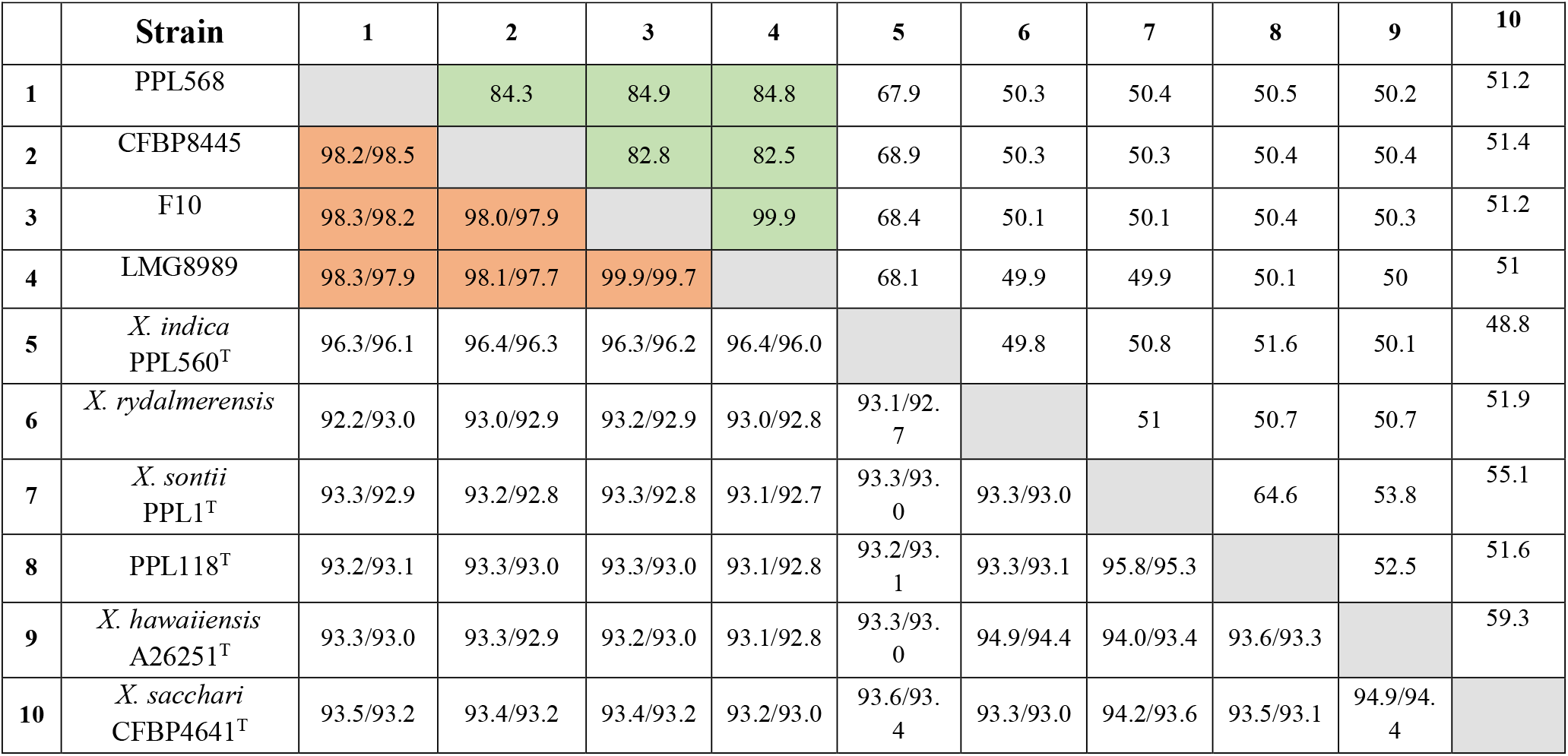
dDDH and orthoANI/ANIb comparisons between *Xanthomonas* strains from this study and type strains of *X. indica, X. sontii, X. sacchari, X. protegens*, and *X. rydalmensis*. Lower triangle shows orthoANI/ANIb values; upper triangle displays dDDH values. Green and light orange shading indicate values above cut-off thresholds for dDDH and ANI, respectively.

To validate the novel species designation, the genomes of strains from this novel lineage were analyzed using the TYGS platform for phylogenetic confirmation (10). TYGS constructed whole-genome phylogenies using query genomes alongside the top closest type strains, delineating species boundaries via colour-coded bars. PPL568 and associated strains clustered within one uniform colour block, indicating shared species and subspecies affiliation **(Fig. 2)**. In contrast, *X. indica* isolates formed separate clusters in distinct colour, confirming species-level divergence.

**Figure 2.**
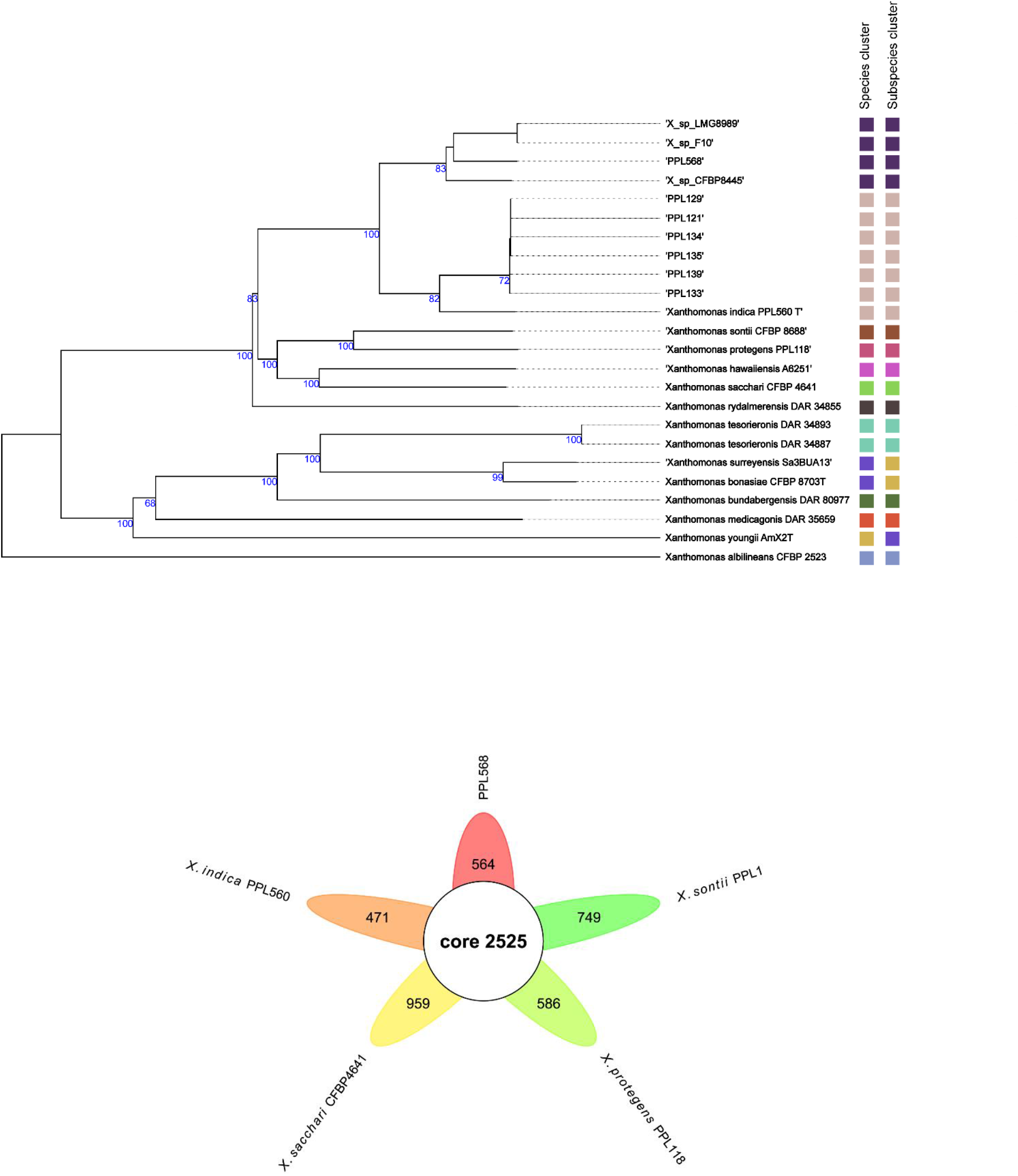
TYGS whole-genome phylogeny of PPL568 and novel lineage isolates (dark blue clade) with top database hits. Branch labels indicate GBDP pseudo-bootstrap support values (>60% shown) from 100 replicates (average 95.5%). Colored bars preceding strain names delineate species clusters and subspecies clusters.

### Comparative Pangenome and COG Analysis

Pangenome analysis revealed that PPL568, a reference strain from the lineage, harbours more unique genes than *X. indica* PPL560^T^, indicating greater genomic flexibility despite overall core genome conservation **(Fig. 3a)**.

**Figure 3.**
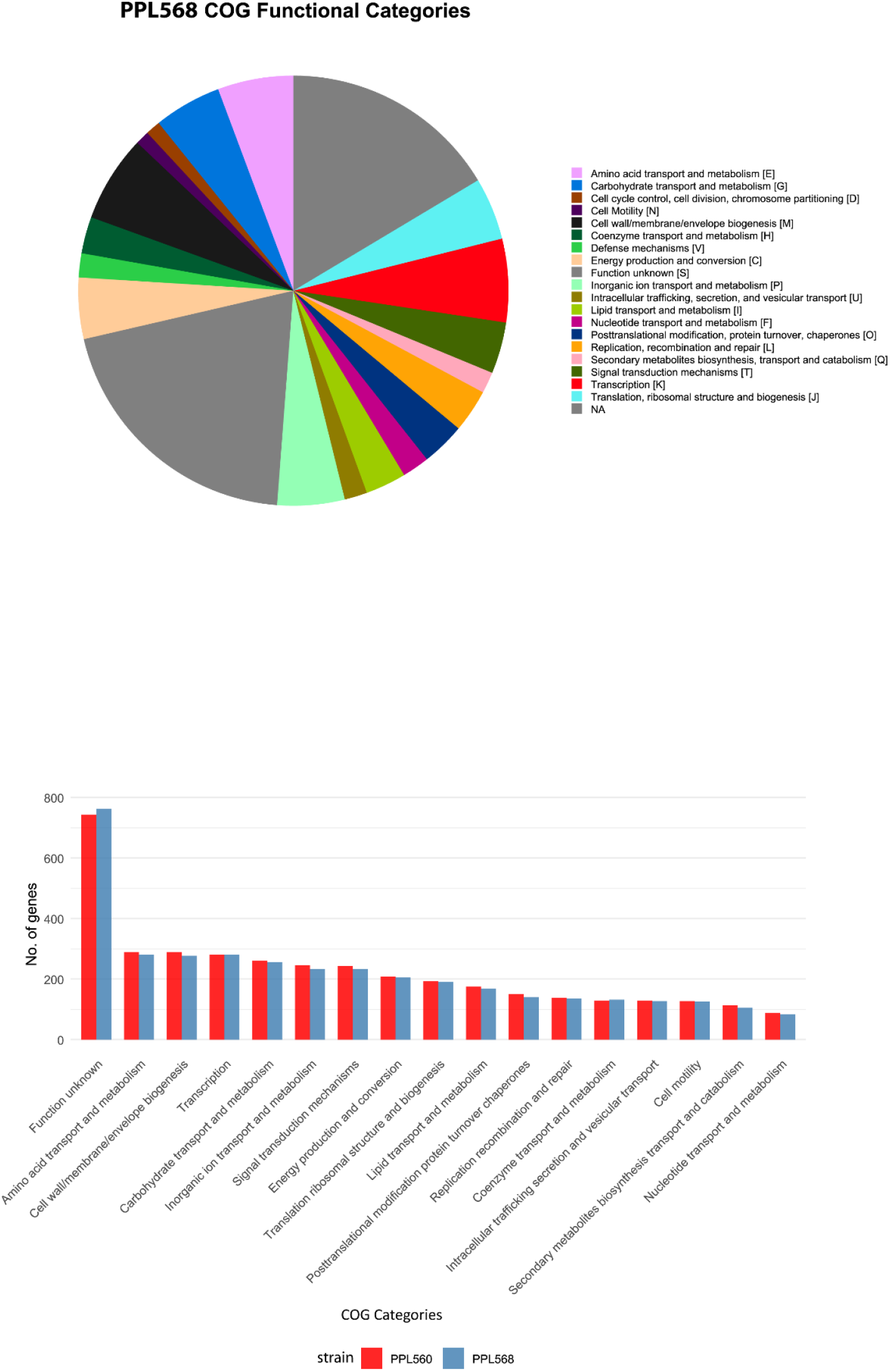
**(a)** Venn diagram of PPL568 and other closely related non-pathogenic *Xanthomonas* species showing core genes at the centre and unique genes in the flower petals. (b) Functional annotation of the PPL568 genome via COG classification. (c) Bar charts compare PPL560^T^ and PPL568 in the percentage distribution of functional categories.

PPL568 were classified into Clusters of Orthologous Groups (COG) categories. About 20.2% of the genome fell into the Function unknown [S] group, with another 16.4% unassigned to any COG class (NA), meaning over 36% remain functionally uncharacterized (Fig. 4a). In the classified functional categories, leading groups for cellular processing and signaling were Signal transduction mechanisms [T] (3.9%), Transcription [K] (6.3%), and Cell wall/membrane/envelope biogenesis [M] (6.5%). Metabolic functions were also well represented, particularly in amino acid [E] (5.7%), carbohydrate [G] (5.1%), and inorganic ion [P] (5.1%) transport and metabolism (Fig. 4). Other vital categories included Translation [J] (4.7%) and Energy production and conversion [C] (4.6%), providing a comprehensive view of the strain’s genetic repertoire **(Fig. 3b)**. The dominance of Function unknown [S] and NA categories highlights a significant reservoir of uncharacterized genes that likely drive the unique phenotypic traits of PPL568. Comparative analysis of COG profiles revealed that, similar to strain PPL560^T^, approximately 35–40% of protein-coding sequences in PPL568 remained unassigned or were categorized under Function Unknown [S], representing an extensive reservoir of uncharacterized proteins typical of bacterial genomes **(Fig. 3c)**. Translation/ribosomal biogenesis [J] dominated both strains (∼25%), followed by amino acid transport/metabolism [E] (∼15-18%) and energy production [C] (∼12%), with PPL560^T^ exhibiting consistently higher absolute gene counts due to its larger genome. The grouped comparison (blue/red side-by-side bars) highlights PPL560^T^ enrichment in amino acid transport/metabolism [E] compared to PPL568, suggesting enhanced nutrient acquisition and regulatory flexibility, while both strains maintain similar relative proportions in cell wall/membrane biogenesis [M] (∼7-8%) and inorganic ion transport/metabolism [P] (∼3-4%), indicating conserved cellular integrity mechanisms. This quantitative distinction in the proportionally conserved COG architecture across strains supports the proposal that PPL568 represents a novel species and confirms its close functional relatedness within *Xanthomonas indica* **(Fig. 3c)**.

**Figure 4.**
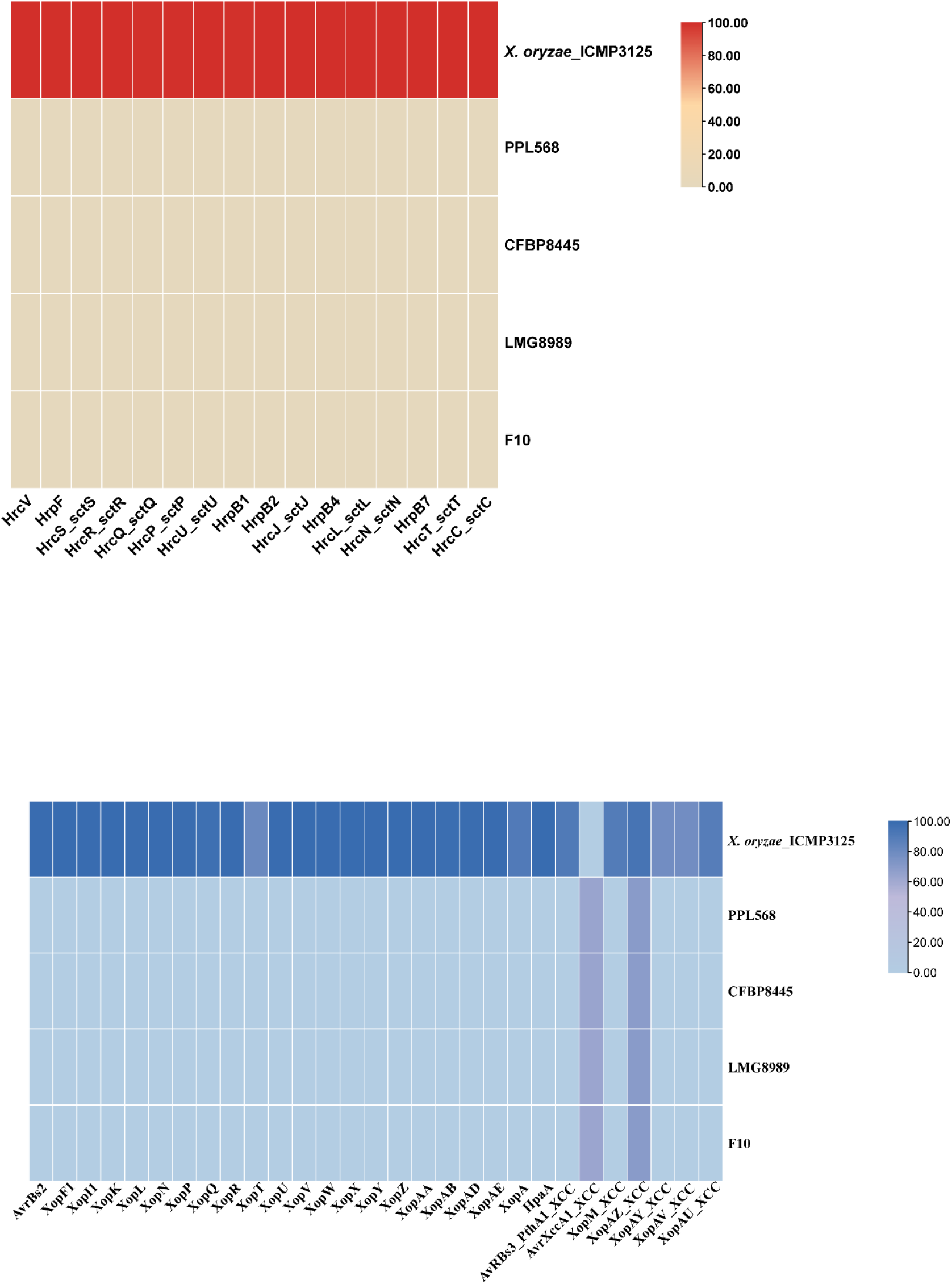
(a) T3SS gene cluster absence in PPL568 and novel lineage compared to *Xanthomonas oryzae* pv. *oryzae*. (b) T3E homolog distribution across xanthomonads, showing a complete repertoire in pathogenic *Xanthomonas oryzae* pv. *oryzae* versus absence in non-pathogenic novel strains

### Lack of Type III Secretion System (T3SS) and Effectors (T3Es)

Genomes of strains from the novel lineage were screened for T3SS and T3E gene homologs, using previously characterized T3SS and T3E repertoires from *Xanthomonas*. Unlike pathogenic species such as *Xanthomonas oryzae* pv. *oryzae*, which carries a complete T3SS, cluster and all associated T3Es, these strains lack T3SS genes and T3E homologs **(Fig. 4a and 4b)**. The non-pathogenic nature of PPL568 and related strains can be explained by the absence of Type III secretion system components, distinguishing this lineage from rice pathogens such as *X. oryzae* pv. *oryzae*. This T3SS deficiency supports their ecological adaptation as commensals or endophytes across diverse hosts (rice, orange, banana), rather than specialized pathogens reliant on effector delivery. Additionally, MacSyFinder analysis of PPL568 identified Type I, Type II, Type IV, and Type V secretion systems, whereas Type VI secretion systems were recognized pathogenicity, factors in *Xanthomonas* were absent. **(Supplementary Table 4)**. Interestingly, PPL568 isolated from rice has also been reported as non-pathogenic, demonstrating a protective effect against the bacterial leaf blight pathogen (6). This lineage is phylogenetically closely related to established non-pathogenic *Xanthomonas* (NPX) species from rice, such as *X. indica* and *X. sontii*.

### Bioprotection potential of NPX strain from the novel lineage

Genome scanning of strains from the novel lineage using established software for antimicrobial locus prediction (27) revealed the presence of ribosomally synthesized and post-translationally modified peptides (RiPPs), atropopeptides, and non-ribosomal independent siderophores **(Fig. 5a)**. These biosynthetic gene clusters suggest a robust chemical arsenal for interbacterial competition and niche exclusion as reported in other NPX species from sub-clade IB (ref-AEM Rekha).

**Figure 5.**
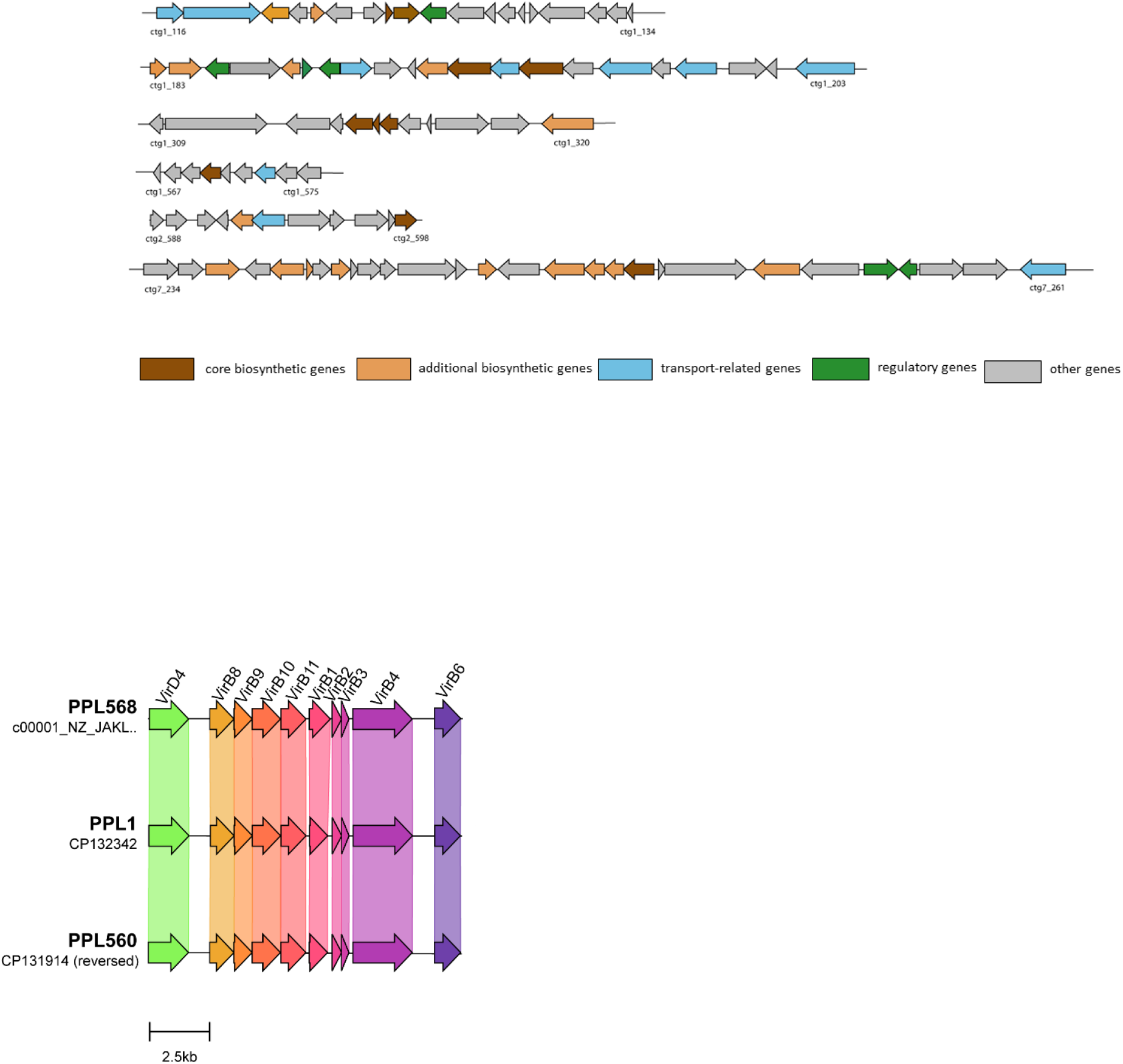
(a) In silico analysis of secondary metabolite biosynthetic gene clusters (BGCs) in the genome of PPL568. (b) Type IV secretion system (X-T4SS) gene cluster for bacterial killing. Homologous genes across strains are shown as identically colored arrows. Gene names appear above arrows.

Prior research has shown that the non-pathogenic strain PPL568 protects rice from bacterial leaf blight caused by the *Xanthomonas oryzae* pv. *oryzae* (Xoo)through multifactorial bioprotection (6). This aligns with the identified antimicrobial loci, suggesting that PPL568 is a promising biocontrol agent. *Xanthomonas* Type IV secretion system (X-T4SS) synteny and genomic architecture in PPL568 are significantly preserved with those in *X. indica* and *X. sontii* **(Fig. 5b)**. Given T4SS substrates are frequently correlated with bacterial killing and interbacterial antagonism (26, 27), this system likely enables NPX strains to competitively exclude the pathogen Xoo from the rice.

### Phenotypic Characterization of strain PPL568

Biochemical profiling using the Biolog GEN III MicroPlate assay assessed carbon source utilization, chemical sensitivity, and growth under varying osmotic and pH conditions across 95 substrates. PPL568^T^ utilized α-D-glucose, D-fructose, D-galactose, D-mannose, dextrin, D-maltose, sucrose, D-trehalose, D-cellobiose, gentiobiose, α-D-lactose, D-turanose, D-melibiose, β-methyl-D-glucoside, D-salicin, N-acetyl-D-glucosamine, N-acetyl-β-D-mannosamine, gelatin, L-glutamic acid, L-alanine, L-serine, L-aspartic acid, formic acid, citric acid, L-lactic acid, acetic acid, propionic acid, bromo-succinic acid, methyl pyruvate, and Tween 40. The strain tolerated pH 5-6 and grew in 1-4% NaCl, 1% sodium lactate, glycerol, and lithium chloride, while resisting lincomycin, rifamycin SV, tetrazolium violet, and vancomycin. Full biochemical details are shown in (**Supplementary Table 2)**. PPL568 and PPL560^T^ exhibited highly similar carbon utilisation patterns, differing only in the assimilation of quinic acid, lithium chloride, and formic acid **(Supplementary Table 3)**. These minor metabolic differences in (formic acid and lithium chloride utilization) distinguish the PPL568 lineage from *X. indica* despite genomic proximity, supporting species-level separation while confirming close relatedness.

## Conclusion

This study establishes PPL568 and closely related strains (F10, LMG8989, CFBP8445, and NCPPB1131) as a novel *Xanthomonas* species through comprehensive genomic and phenotypic characterization. Core-genome phylogeny, TYGS analysis, and dDDH values confirm the novel lineage’s monophyletic clustering distinct from *X. indica*, exceeding species-level thresholds internally while remaining below *X. indica* cutoffs. Rice seed non-pathogenic *Xanthomonas* (NPX) form a distinct species complex exhibiting unprecedented host adaptability across rice, orange, and banana. The absence of T3SS/T3E genes explains non-pathogenic endophytic behaviour. Phylogenomic, pangenomic, and taxono-genomic analyses provide robust evidence delineating this lineage from *X. indica* while positioning it as a close relative. These findings support the proposal of *Xanthomonas imtechensis* sp. nov. (Type strain PPL568^T^), highlighting NPX diversity and probiotic potential in rice microbiomes.

### Description of *Xanthomonas imtechensis* sp. nov

*Xanthomonas imtechensis* (im.tech.en’sis. N.L. masc. adj. imtechensis, in with respect to IMTECH, the Institute of Microbial Technology) is a novel species within the genus *Xanthomonas*.

PPL568^T^ obtained from rice seeds of healthy plants, is rod-shaped, Gram-negative, and grows aerobically. It forms convex, mucoid, pale-yellow colonies on PSA following 48 h at 28 °C. This novel strain catabolized D-galactose, α-D-glucose, D-fructose, dextrin, D-mannose, sucrose, gentiobiose, D-trehalose, β-methyl-D-glucoside, D-salicin, D-melibiose, D-turanose, D-cellobiose, α-D-lactose, N-acetyl-D-glucosamine, N-acetyl-β-D-mannosamine, gelatin, L-serine, L-glutamic acid, L-alanine, L-aspartic acid, citric acid, formic acid, propionic acid, L-lactic acid, acetic acid, bromo-succinic acid, methyl pyruvate, and Tween 40. Growth was supported at pH 5–6 and in presence of 1–4% NaCl, glycerol, lithium chloride, L-fucose, and 1% sodium lactate. PPL568^T^ lacks pathogenicity toward rice plants. The type strain genome is 4.7 Mb and has a 69% G+C content. The Type strain is PPL568^T^ = MTCC 13186 = CFBP 9040 = ICMP 24395.

## Acknowldegements

We acknowledge the Department of Science and Technology - Innovation in Science Pursuit for Inspired Research (DST-INSPIRE) fellowship to AS.

## Author contribution

AS performed genome analysis, and drafted the manuscript. PBP conceptualized the study, contributed to design, and finalized the manuscript.

## Funding

This study received support from the “Microbial Type Culture Collection and Gene Bank (MTCC), serving as a Microbial Resource Centre and International Depository Authority (IDA)” project. It was jointly funded by DBT and CSIR (1:1 ratio) through the DBT-SAHAJ initiative (DBT project No. BT/INF/22/SP43064/2022; CSIR project No. MLP065).

## Conflict of interest

The authors declare no conflicts of interest.

## Notes

### Competing Interest Statement

The authors have declared no competing interest.

