## Supplementary Table 1 for "*Xanthomonas imtechensis* sp. nov. - a novel member of non-pathogenic *Xanthomonas* with bioprotection function from healthy rice seeds"

| **S. No.** | **Strains** | **dDDH** | **ANI** |
| --- | --- | --- | --- |
|  |  | **PPL568^T^** | **PPL568^T^** |
| 1. | CFBP8445 | 84.3 | 98.2 |
| 2. | F10 | 84.9 | 98.3 |
| 3. | LMG8989 | 84.8 | 98.3 |
| 4. | PPL560^T^ | 67.9 | 96.3 |
| 5. | *X. rydalmerensis* | 50.3 | 92.2 |
| 6. | *X. hawaiiensis* A26251^T^ | 50.2 | 93.3 |
| 7. | *X. sacchari* CFBP4641^T^ | 51.2 | 93.5 |
| 8. | *X. protegens* PPL118^T^ | 50.5 | 93.2 |
| 9. | *X. sontii* PPL1^T^ | 50.4 | 93.3 |

Digital DNA–DNA hybridization (dDDH), average nucleotide identity (ANI) values for strain PPL568^T^ were determined by comparison with type and representative strains of other *Xanthomonas* species.
