## Supplementary Table 2 for "*Xanthomonas imtechensis* sp. nov. - a novel member of non-pathogenic *Xanthomonas* with bioprotection function from healthy rice seeds"

| Negative Control | Dextrin | D-Maltose | D-Trehalose | D-Cellobiose | Gentiobiose | Sucrose | D-Turanose | Stachyose | Positive Control | pH 6 | pH 5 |
| --- | --- | --- | --- | --- | --- | --- | --- | --- | --- | --- | --- |
| D-Raffinose | α-D-Lactose | D-Melibiose | β-Methyl-D-Glucoside | D-Salicin | N-Acetyl-D-Glucosamine | N-Acetyl-β-D-Mannosamine | N-Acetyl-D-Galactosamine | N-Acetyl-Neuraminic Acid | 1% NaCl | 4% NaCl | 8% NaCl |
| D-Glucose | D-Mannose | D-Fructose | D-Galactose | 3-Methyl Glucose | D-Fucose | L-Fucose | L-Rhamnose | Inosine | 1% Sodium Lactate | Fusidic Acid | D-Serine |
| D-Sorbitol | D-Mannitol | D-Arabitol | Myo-Inositol | Glycerol | D-Glucose-6-PO4 | D-Fructose-6-PO4 | D-Aspartic Acid | D-Serine | Troleandomycin | Rifamycin SV | Minocycline |
| Gelatin | Glycyl-L-Proline | L-Alanine | L-Arginine | L-Aspartic Acid | L-Glutamic Acid | L-Histidine | L-Pyroglutamic Acid | L-Serine | Lincomycin | Guanidine HCl | Niaproof 4 |
| Pectin | D-Galacturonic Acid | L-Galactonic Acid Lactone | D-Gluconic Acid | D-Glucuronic Acid | Glucuronamide | Mucic Acid | Quinic Acid | D-Saccharic Acid | Vancomycin | Tetrazolium Violet | Tetrazolium Blue |
| p-HydroxyPhenylacetic Acid | Methyl Pyruvate | D-Lactic Acid Methyl Ester | L-Lactic Acid | Citric Acid | α-Keto-Glutaric Acid | D-Malic Acid | L-Malic Acid | Bromo-Succinic Acid | Nalidixic Acid | Lithium Chloride | Potassium Tellurite |
| Tween 40 | γ-Amino-Butryric Acid | α-HydroxyButyric Acid | β-Hydroxy-D,Lbutyric Acid | α-Keto-Butyric Acid | Acetoacetic Acid | Propionic Acid | Acetic Acid | Formic Acid | Aztreonam | Sodium Butyrate | Sodium Bromate |

**Supplementary Table : 2** The biochemical profile of PPL568 was determined using a BIOLOG GEN III microplate, which includes assays for utilization of diverse carbon sources, resistance to various antibiotics, and growth at different pH and NaCl concentrations. A Grey-shaded cell denotes a positive reaction, a red-shaded cell indicates a negative reaction, and a yellow-shaded cell represents a variable reaction.

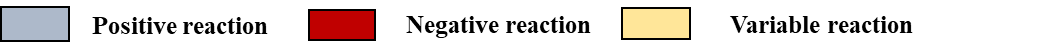
