## Supplementary Table 3 for "*Xanthomonas imtechensis* sp. nov. - a novel member of non-pathogenic *Xanthomonas* with bioprotection function from healthy rice seeds"

**Supplementary Table: 3**	Phenotypic profiling of	PPL568 compared with *X.indica* PPL560^T*	(*^ Biolog data taken from literature). Symbols indicate source utilization (+) or lack of metabolism (-), with "v" denoting variable readings.

| **S. No.** | **BIOLOG tests** | **PPL568^T^** | ***X. indica* PPL560^T^** |
| --- | --- | --- | --- |
| **1** | Dextrin | + | + |
| **2** | D-maltose | + | + |
| **3** | D-Trehalose | + | + |
| **4** | D-Cellobiose | + | + |
| **5** | Gentiobiose | + | + |
| **6** | Sucrose | + | + |
| **7** | D-turanose | + | + |
| **8** | D-raffinose | - | - |
| **9** | α-D-lactose | + | + |
| **10** | D-melibiose | + | + |
| **11** | β-methyl-D-glucoside | + | + |
| **12** | Gelatin | + | + |
| **13** | D-fructose | + | + |
| **14** | D-galactose | + | + |
| **15** | L-fucose | + | v |
| **16** | L-rhamnose | - | - |
| **17** | D-sorbitol | - | - |
| **18** | D-mannitol | - | - |
| **19** | D-arabitol | - | - |
| **20** | L-alanine | + | + |
| **21** | L-aspartic acid | + | + |
| **22** | L-glutamic acid | + | + |
| **23** | Formic acid | + | - |
| **24** | Quinic acid | v | + |
| **25** | Citric acid | + | + |
| **26** | Bromo-succinic acid | + | + |
| **27** | Propionic acid | + | + |
| **28** | Acetic acid | + | + |
| **29** | α-D-Glucose | + | + |
| **30** | D-Mannose | + | + |
| **31** | L-Arginine | - | - |
| **32** | L-Histidine | - | v |
| **33** | D-Saccharic Acid | - | - |
| **34** | L-Lactic Acid | + | + |
| **35** | Lithium chloride | + | - |
